# Mutational effects on carbapenem hydrolysis of YEM-1, a new sub-class B2 metallo-β-lactamase from *Yersinia mollaretii*

**DOI:** 10.1101/2020.01.29.926220

**Authors:** P. S. Mercuri, R. Esposito, S. Blétard, S. Di Costanzo, M. Perilli, F. Kerff, M. Galleni

## Abstract

The analysis of the genome sequence of *Yersinia mollaretii* (*Y. mollaretii*) ATCC 43969 indicates the presence of the *bla_YEM_* gene coding for YEM-1, a putative subclass B2 metallo-β-lactamase. The objectives of our work were to produce, purify and complete the kinetic characterization of YEM-1. Compared to the known subclass B2 metallo–β-lactamases, YEM-1 displayed a narrowest substrate profile since it is only able to hydrolyse imipenem with a high catalytic efficiency but not all the other carbapenems tested such as biapenem, meropenem, doripenem and ertapenem. A possible explanation of this peculiar activity profile is the presence of tyrosine 67 (loop L1), threonine 156 (loop L2) and serine 236 (loop L3) respectively. We showed that the substitution of Y67 broadened the activity profile of the enzyme for all carbapenems but still displayed a poor activity toward the other β-lactam classes.

## INTRODUCTION

The genus *Yersinia* belonging to the family of *Enterobacteriaceae* shows three mains groups of human pathogen strains, *Yersinia pestis* (*Y. pestis*)*, Yersinia pseudotuberculosis* (*Y. pseudotuberculosis*) and *Yersinia enterolitica* (*Y. enterocolitica). Y. pestis*, the bacterial agent of bubonic plague, escapes the macrophages action and is the cause of a lethal bacteremia without antibiotic treatments (1). A new threat is the emergence of a multidrug resistant *Y. pestis* strain (2). *Y. pseudotubercolis* and *Y. enterocolitica* are important cause of gastroenteritis in human following the ingestion of undercooked or contaminated pork and vegetables (3–4). Others *Yersinia* species like *Y. fredricksenii, Y. kristensenii, Y. intermedia, Y. mollaretii, Y. bercovieri and Y. rohdei* are considered as not pathogen for human (5–6). Our interest was focused on *Y. mollaretii*. This specie, previously known as *Y. enterocolitica*-like organism biogroup A, was renamed *Yersinia mollaretii* and assigned in the new biogroup 3A by Wauters et al (7). The majority of those strains were isolated from environmental sources such as soil, drinking water, raw vegetables, and meat but few were clinical isolates from patient stools affected by gastrointestinal disease. It was demonstrated that it can colonize the low level of the human ileum but the absence of human virulence markers classified this strain as non-pathogenic and saprophyte (8). The sequence genome of the reference strain *Y. mollaretii ATCC 43969* (numbered as CCUG26331, CDC2465-87, CIP103324 and DSM18520 in other biobanks) was determined (NCBI Reference Sequence: NZ_AALD02000006.1). Its analysis indicated the presence of two β-lactamase genes, namely *blaB* coding for an AmpC-like enzyme and one coding for a putative sub–class B2 metallo β-lactamase called YEM-1 (**YE**rsinia **M**etallo-β-lactamase-GenBank: EEQ11779.1). It is interesting to note the absence of the *blaA* gene encoding for an Extented Spectrum class A β-Lactamase found in other *Yersinia* species (9–12). This observation is in good agreement with the phenotypic studies showing that *Y. mollaretii* displayed a resistant pattern to amoxillin and an intermediate pattern of susceptibility to amoxicillin/clavulanic acid and cefoxitin (13). These studies demonstrated a clear relationship between the expression of BlaA and BlaB by *Yersinia* strains and their antibiotic resistance patterns. Up to date, the impact of the production of YEM-1 on the resistance pattern is unknown. Compared to the characterized sub-class B2 MBLs (14), such as CphA (15), SfhI (16) and ImiS (17), the amino acids involved in the two zinc binding sites were strictly conserved, (N116, H118 and H196 for Zn 1 and D120, C221 and H263 for Zn2). CphA is the most studied enzyme of the sub-class B2 MBLs (18–20). It is a strict carbapenemase and the binding of a second zinc ion inhibited its activity (21). A model of the three-dimensional structure of YEM-1 was build using the crystallographic structure of CphA as template (Figure 1). As expected, the two enzymes possessed a similar fold. The main difference between the two enzymes was the substitution V67Y in YEM-1 (22). V67 is conserved in all known sub-class B2 and also in members of sub-class B1 such as BcII, IMP-1, CcrA and NDM-1 but not in VIM-1 where the same substitution V67Y was observed. This residue is part of L1 loop and is involved in the hydrophobic wall of the active site that plays an important role in the carbapenem binding especially for biapenem (23). Compared to CphA and the other known sub-class B2 MBLs, two other residues (F156 and F236) are not conserved in YEM-1. In CphA, these residues are involved in an interaction between the biapenem intermediate and the enzyme (24). Moreover, F236 is a part of a mobile loop L3, localised at the entrance of the active site (25). In YEM-1, the residues in position 156 and 236 are substituted by a threonine and a serine respectively.

**FIGURE 1.**
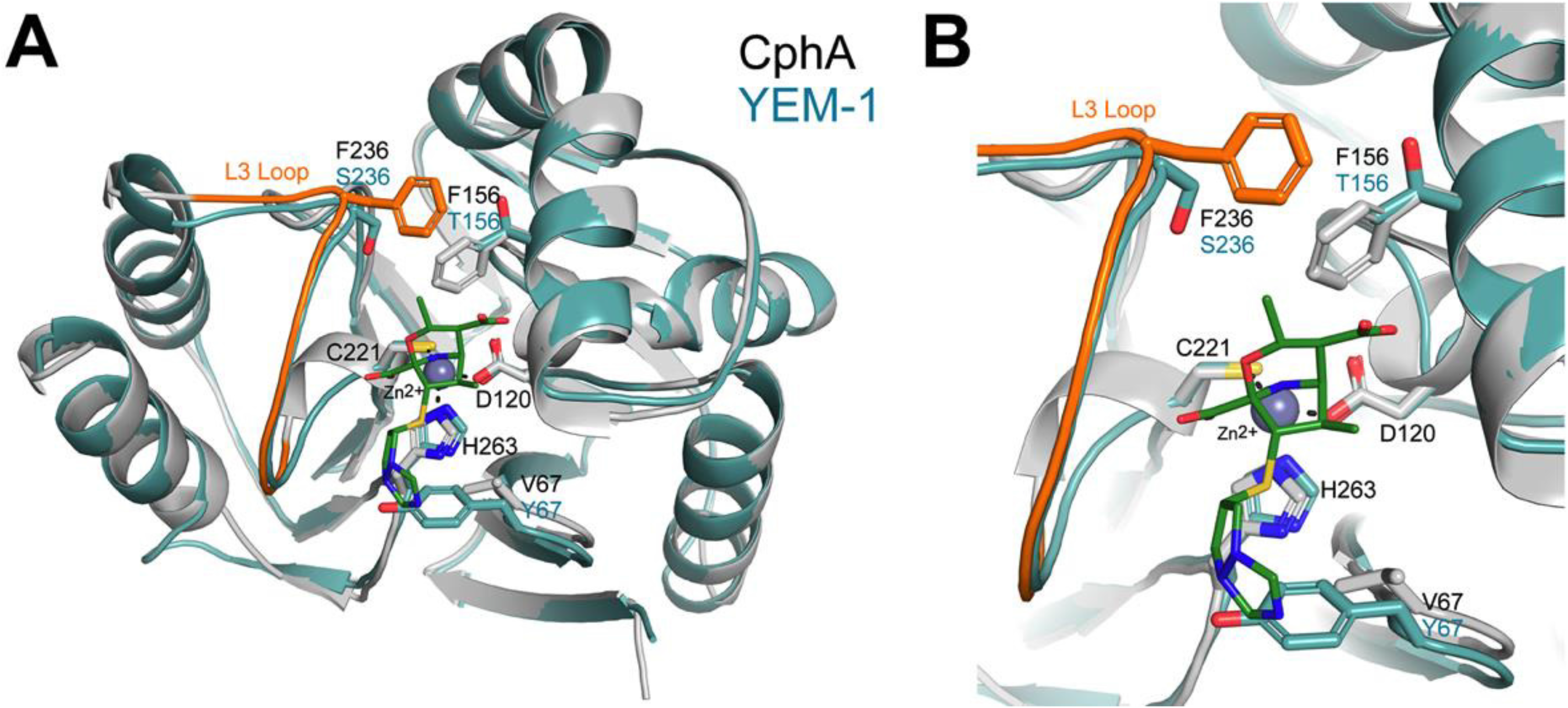
A cartoon representation of the CphA structure (grey) in complex with biapenem (green sticks), superposed to the homology model of YEM-1 (blue). The zinc ion present in the active site is displayed as a grey sphere. **B.** The residues coordinating the zinc ion as well as residues in postion 67, 156 and 236 are showed as sticks.

The aims of this work were: i) to characterize the kinetic properties of the wild-type YEM-1; ii) to evaluate the impact of the substitution of Y67, T156 and S236 on its catalytic efficiency by the study of the different single mutants, the double mutant T156F/S236F and the triple mutant Y67V/T156F/S236F.

## RESULTS

The *bla_YEM_* gene (747 bp) was amplified by PCR using the *Y. mollaretii* ATCC 43969 genomic DNA as template. After the verification of the correct nucleotidic sequence of the amplimer, the gene was sub-cloned in pET26b for enzyme expression. The *bla_YEM_* contains 15 rare codons that could affect the expression of the enzyme in *Escherichia coli* (*E. coli*). Therefore, we decided to use the *E. coli* Rosetta^TM^ (DE3) strain as host to be able to improve the protein production yield. A large quantity of protein was produce as cytoplasmic inclusion bodies when the cultures were grown at 37°C. We succeeded to produce a soluble form of YEM-1 when the cultures were realized O/N at 18°C in presence of IPTG 0.1 mM (final concentration). The enzyme was purified in three chromatographic steps as described in the section “Material & Methods” with a final yield of 6 mg/L of culture. The analysis of the protein solution by ESI-Q-TOF mass spectrometry confirmed the homogeneity of the protein and measured the molecular weight of the mature form of the β-lactamase. Its value (25634 Da) indicated that the pre-β-lactamase include a signal peptide of 20 amino acids and the mature enzyme contains 229 amino acids with the N-terminal sequence corresponding to NH2-SKIA. A blast-P analysis of the known *Yersinia* genomes showed the presence of the YEM-1-like MBL in other *Y. mollaretii* strains but also in *Y. intermedia* and *Yersinia massiliensis* strains. A clustalW aligment of these enzymes showed a minimal identity value of 90% (Figure 2). We noted that Y67 and S236 are strictly conserved in all the YEM-like sequences. We made the hypothesis that the only significant difference between the *Y. mollaretii*, *Y. intermedia* and *Y. massiliensis* is the presence of F156. YEM-1 hydrolysed carbapenems but we could not detect any hydrolytic activities toward penicillins or cephalosporins. These preliminary data confirmed that YEM-1 belongs to the subclass B2 MBLs. Therefore, we were only able to calculate the steady state kinetic parameters against a representative set of carbapenem antibiotics (Table 1). Compared to CphA, YEM-1 efficiently hydrolysed only imipenem. For the others carbapenems, we noted a large decrease of the k_cat_ value and a major increase of K_m_. All these data confirmed that YEM-1 possesses a narrow activity spectrum. For example, its catalytic efficiencies against doripenem and biapenem are 100 and 430 fold lower compared to imipenem (Table 1). We can conclude that YEM-1 is an imipenemase. We expected that, as observed for the other members of the subclass B2 MBL, the activity of YEM-1 is maximal for the mono-zinc form. We showed that its activity is affected by the presence of increasing concentration of Zn^2+^, Co^2+^ and Cd^2+^ (Figure 3 A-C). Interestingly, YEM-1 displayed a lower affinity (apparent K_d_ = 360 μM) for the Zn2 compared to CphA (K_d_ = 46 μM) and its residual activity in presence of 4mM Zn^2+^ is equal to 40 % of its activity at 1 μM Zn^2+^. In the case of Co^2+^ and Cd^2+^ ions, we observed a complete inactivation of the wild-type and mutant enzymes at 1 mM metal ion concentration. This phenomenon may be due to the substitution of Zn1 and /or the binding of three metal ions. The production and purification of five YEM-1mutants (three single mutants Y67V, T156F, S236F, the double mutant T156F-S236F and the triple mutant Y67V-T156F-S236F) were realised in the same conditions defined for the wild-type enzyme. The kinetic results confirm that, together, all the mutations do not favour the broadening of the activity profile of YEM against other β-lactam classes neither its apparent affinity for the binding of the second zinc ion (Table 1). The substitution of Y67V yielded a global increase of the catalytic efficiency to carbapenems, with the exception of imipenem. It is interesting to note that the mutant YEM-1Y67V has an activity profile similar to CphA. For ertapenem and meropenem, the values of K_m_ are lower than that of the YEM-1 WT and comparable to CphA. The mutations of T156 and S236 in phenylalanine decrease the K_m_ value for imipenem compared to YEM-1WT and CphA but the K_m_ for other tested carbapenems were not modified. It is to be noted that all the catalytic efficiency of the two mutants decreases compared to the WT enzyme. The characterization of triple mutant Y67V T156F S236F mutant highlighted that the presence of a valine residue in position 67 is essential to favour a broader activity profile toward carbapenems (Table 1). With the exception for doripenem, the results showed that the mutation Y67V provides at a better activity towards the carbapenems. For T156F, S236F and the double mutant T156F-S236F, we noted that the mutations affected the K_m_ values and by consequence, the catalytic efficiency of the mutants compared to YEM-1 with the exception of imipenem.

**FIGURE 2.**
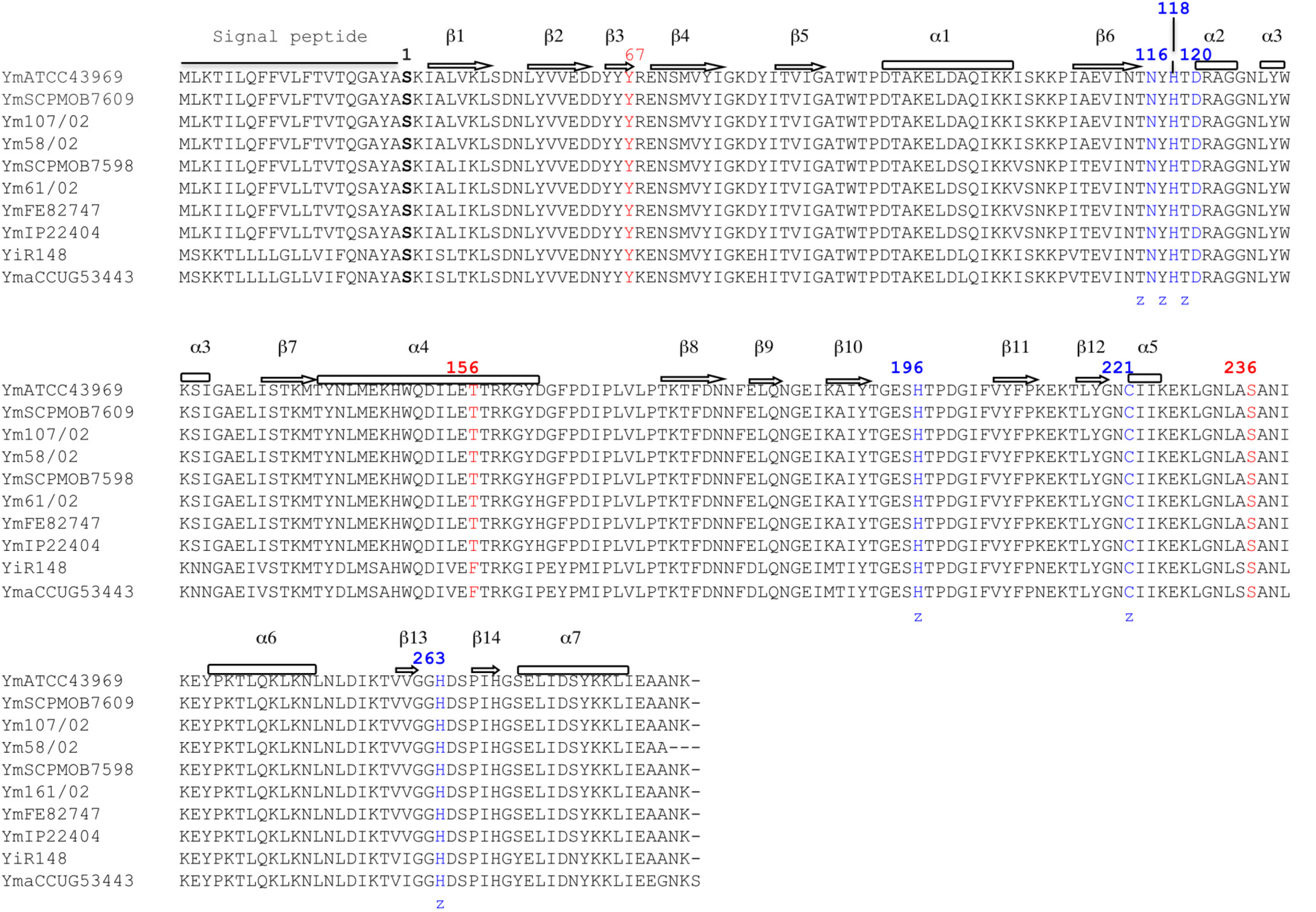
ClustalW alignment of subclass B2 metallo-β-lactamase produced by *Yersinia* species. Ym (*Yersinia mollaretii*) ATCC43969 (WP_004874166.1), Ym SCPM-O-B-7609 (WP_ 004874166.1), Ym 107/02 WP_004874166.1, Ym 58/02 (WP_050540509.1), Ym SCPM-O-B-7598 (WP_049645673.1), Ym 61/02 (WP_049645673.1), Ym FE82747 (WP_049645673.1), Ym P22404 (WP_049645673.1), Yi (*Yersinia intermedia*) R148 (WP_019211763.1) and Yma (*Yersinia massiliensis*) CCUG 53443 (WP_019211763.1). Residues at position 67, 156 and 236 are in red (BBL numbering). The zinc ligands residues are in blue.

**TABLE 1.** Kinetic parameters of YEM-1WT and mutants compared to CphA. Kd are the apparent dissociation constants values for zinc ions. The numbers in parenthesis correspond to the values of the ratios of (kcat/Km)Imipenem/ (kcat/Km)carbapenem for each protein.

| Antibiotics | Kinetic parameters | CphA | YEM-1 | Y67V | T156F | S236F | T156F-S236F | Y67V-T156F-S236F |
| --- | --- | --- | --- | --- | --- | --- | --- | --- |
| **Imipenem** | k<sub>cat</sub> (sec<sup>-1</sup>) | 1320 | 480 | 1770 | 190 | 170 | 760 | 40 |
|  | K<sub>m</sub> (μM) | 310 | 160 | 270 | 32 | 90 | 20 | 60 |
|  | k<sub>cat</sub>/K<sub>m</sub> (μM<sup>-1</sup> sec<sup>-1</sup>) | **3.2 (1)** | **3 (1)** | **7 (1)** | **6 (1)** | **2 (1)** | **38 (1)** | **0.7 (1)** |
| **Meropenem** | k<sub>cat</sub> (sec<sup>-1</sup>) | 1440 | >400 | 800 | >230 | >15 | >75 | >3300 |
|  | K<sub>m</sub> (μM) | 630 | >2500 | 400 | >2500 | >2500 | >2500 | >2500 |
|  | k<sub>cat</sub>/K<sub>m</sub> (μM<sup>-1</sup> sec<sup>-1</sup>) | 2.23 (1.4) | 0.15 (20) | **2 (3.5)** | 9 10<sup>-2</sup> (66) | 10<sup>-2</sup> (200) | 3 10<sup>-2</sup> (1260) | **1.3** |
| **Biapenem** | k<sub>cat</sub> (sec<sup>-1</sup>) | 1420 | >16 | >1000 | >11 | >0.8 | >15 | >300 |
|  | K<sub>m</sub> (μM) | 1500 | >2500 | >2500 | >2500 | >2500 | >2500 | >2500 |
|  | k<sub>cat</sub>/K<sub>m</sub> (μM<sup>-1</sup> sec<sup>-1</sup>) | 0.96 (3) | 7 10<sup>-3</sup> (430) | **0.4 (17)** | 4 10<sup>-3</sup> (1500) | 3 10<sup>-4</sup> (6.6 10<sup>4</sup>) | 0.006 (6300) | **0.13 (6)** |
| **Ertapenem** | k<sub>cat</sub> (sec<sup>-1</sup>) | 1410 | >400 | 1650 | >400 | >20 | >400 | 810 |
|  | K<sub>m</sub> (μM) | 230 | >2500 | 120 | >2500 | >2500 | >2500 | 80 |
|  | k<sub>cat</sub>/K<sub>m</sub> (μM<sup>-1</sup> sec<sup>-1</sup>) | 6.8 (0.4) | 0.15 (20) | **14 (0.5)** | 0.15 (40) | 0.01 (200) | 0.15 (250) | **8 (1.1)** |
| **Doripenem** | k<sub>cat</sub> (sec<sup>-1</sup>) | 940 | >100 | >1300 | >20 | >5 | >10 | >50 |
|  | K<sub>m</sub> (μM) | 580 | >2500 | >2500 | >2500 | >2500 | >2500 | >2500 |
|  | k<sub>cat</sub>/K<sub>m</sub> (μM<sup>-1</sup> sec<sup>-1</sup>) | 1.6 (2) | 0.03 (100) | **0.5 (14)** | 0.009 (660) | 2 10<sup>-3</sup> (10<sup>3</sup>) | 0.004 (9500) | 0.02 (35) |
|  | *K<sub>d</sub>'(μM) Zn2* | *46* | *360* | *330* | *410* | *300* | *400* | *270* |

Finally, all the enzymes tested were inhibited in the presence of zinc ion. The apparent Kd values for the binding of the second zinc are comparable between the WT and the mutants (Table 1). The mutations did not affect the influence of metal ions on the activity of YEM-1 nor their binding of Co^2+^ and Cd^2+^.

The YEM-1 model obtained with Yasara is very similar to the structure of CphA used as template (rmsd of 0.36 Å over 209 Cα) and a single 3 amino acids insertion before the C-terminal helix that is not expected to affect substrate hydrolysis. The active site is also well conserved, with a rmsd of 0.32 Å for all atoms of the six residues involved in zinc binding. Compared to CphA, YEM-1 presents three significant differences (Garau 2005), a tyrosine at the position 67 would interfere with the bulky side chain of this the active site surrounding: the replacement of V67 by a tyrosine and the substitution of F156 and F236 by a threonine and a serine respectively. According to the CphA:biapenem complex (pdb code 1X8I carbapenem). On the other side of the active site, the two other mutations remove the hydrophobic interactions between the two phenylalanines present in CphA and widen the catalytic site cavity.

## DISCUSSION

Despite the high percentage of sequence identity (58 %) between YEM-1 and CphA, we noted that YEM-1 is less effective than CphA against carbapenems. With the exception of imipenem, the catalytic efficiency of YEM-1 decreases at least a factor 10. Furthermore, for the majority of carbapenems, it was not possible to determine the individual kinetic constants because the K_m_ are higher than the substrate concentrations tested yielding pseudo-first order kinetics. The analysis of the structure of the complex CphA-hydrolysed biapenem, showed the importance of H263 in the interaction with biapenem sulphur atom and of V67 and W83 to stabilize the carbapenem intermediate in the active site (24). H263 and W87 are conserved in YEM-1 but it contains a tyrosine in position 67 instead of a valine as observed for all the other subclass B2 and the majority of subclass B1 MBLs. The importance of Val in position 67 is confirmed by the higher ability of the mutant Y67V to hydrolysis carbapenems. A computational analysis on the generation of CphA variants under imipenem selection (26) shows that position 67 is tolerant to multiple amino acid substitutions. The different mutants retained a catalytic efficiency comparable to the WT enzyme. In the case of YEM-1, we showed that the presence of Y67 reduced its activity towards the other tested carbapenems. Its substitution to valine increased the YEM-1 activity to the levels observed for the other sub-class B2 MBLs. Like CphA, YEM-1 is inhibited in presence of Zn^2+^ ions. Nevertheless, the apparent dissociation constant for zinc is higher for YEM-1 compared to CphA. This result indicates that YEM-1 displayed its maximum of activity in the mono-zinc form and that it is less sensitive that CphA and ImiS in presence of increasing concentration of zinc ion. CphA and ImiS can bind up to 2 equivalent of Zn (II) or Co (II) but the maximal activity is with one equivalent of metal ion (27). The further verification of the higher dissociation constant of YEM-1 for Zn^+2^ would require the production the YEM-1 apoenzyme form and determine the affinity constant for Zn1 and Zn2. (28).

The lower affinity of the inhibitory Zn^2+^ binding site of YEM-1 could result from the F156T and F236S substitutions because they induce the loss of a hydrophobic rigidifying interaction between the two sides of the active site in the vicinity of the second Zn^2+^ binding site (Figure 3 A). We can also conclude that the *Y. intermedia* and *Y. massiliensis* MBLs display a lower catalytic efficiency compared to YEM-1.

**Figure 3.**
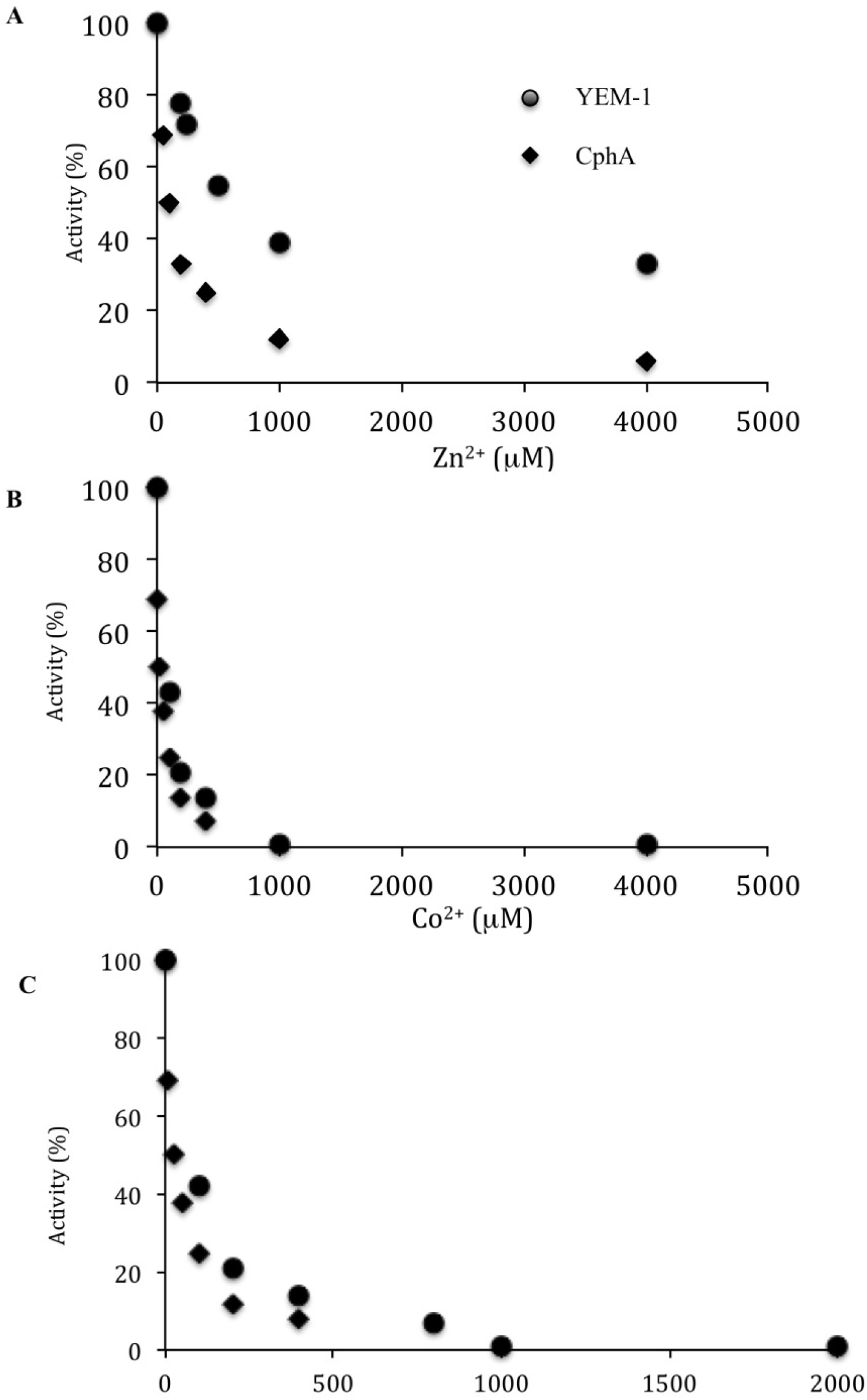
Influence of Zn^2+^ (A), Co^2+^ (B) and Co^2+^ (C) concentrations on CphA and YEM-1 carbapenemase activity

We also noted that the YEM-1 is still active at high zinc ion concentrations. It could be interesting to determine the X-ray structure of the di-zinc enzyme in order to identify the structural determinants that favour the activity of the di zinc MBL.

In conclusion, *Y. mollaretii* produces two β-lactamases potentially able to exhibit a resistance profile covering a large panel of β-lactam antibiotics. In 2017, Ribeiro et al (29) showed the presence of *Y. mollaretii* in the biofilm composition isolated in children dental carie. Therefore, the presence of β-lactamase genes in the opportunist strain *Y. mollaretii* should be not underestimated.

## MATERIALS & METHODS

### Antibiotics, bacterial strains and plasmids

Kanamycin was purchased from MP Biomedicals, France. Ampicillin from SA Bristol-Myers Squibb Belgium,N.V. Imipenem and Ertapenem from MSD, Belgium. Meropenem from Astro Zeneca Belgium. Doripenem and Biapenem were purchased from Sigma Aldrich, Belgium. *E.coli* DH5α (Life Technologies, Belgium) and *E. coli* Rosetta (DE3) (Novagen Inc., Madison, Wis.) were used as recipient on the cloning gene step and enzyme expression. The gene was cloned initially into pJET1.2 vector (Thermo Fischer Scientific, Belgium) and after subcloned into pET-26b (Novagen Inc., Madison, Wis).

### Cloning of bla_YEM-1_ gene

The *bla_YEM-1_* gene was isolated by PCR from the purified genomic DNA of *Y. mollaretii* Wauters DMS18520 (*Y. mollaretii* ATCC43969) purchased from the Leibniz Institute DSMZ (German Collection of Microorganisms and Cell Cultures). The extraction of genomic DNA was realised with the help of Wizard Genomic DNA purification kit (Promega Benelux B.V.). Two oligonucleotides YblaUP_NdeI and YblaRP _BamHI (Table S1) were synthetized on the basis of the sequence of the genome *Y. mollaretii* (NCBI Reference: NZ_AALD02000006.1). PCR conditions were as follows: incubation at 98°C for 30 sec and 35 cycles of amplification (denaturation at 98°C for 10 sec, annealing at 50°C for 30 sec, and extension at 72°C for 30 sec). The Q5Hight-Fidelity DNA polymerase (New England BioLabs Inc.) was used in this work. The PCR product was cloned into the pJET1.2 vector. After the verification of the nucleotidic sequence, *bla*_YEM-1_ was subcloned into pET26b previously digested by the *Nde*I and *Bam*HI restriction enzymes.

### Site -directed mutagenesis

In order to replace the codon for Tyr into the codon for Val in position 67, the oligonucleotides Y67V_fwd and Y67_rev were synthesised (Table S1). The sequences of the oligonucleotides designed to generate the T156F (T156F_fwd and T156F_rev) and S236F (S236F_fwd and S236F _rev) mutants are shown in Table S1. The pET26b-*bla_YEM_* was used as template for the mutagenesis and the amplification conditions were: incubation at 98°C for 30 sec and 30 cycles of amplification (denaturation at 98°C for 10 sec, annealing at 55°C for 5 min, and extension at 72°C for 30 sec). After amplification, the PCR fragments were digested by the *Dpn*I restriction enzyme in order to eliminate the template’s DNA. The double and triple mutants T156FS236F and Y67VT156FS236F were obtained by consecutive PCR. The recombinant plasmids were selected on LB agar plates supplemented with kanamycin 50 μg/mL. The presence of the desired mutations was confirmed, by sequencing.

### Expression and purification of wild-type and mutants of YEM-1

All the enzymes were produced in *E. coli* Rosetta (DE3) carrying pET26b/*bla_YEM-1-_* and pET26b/*bla_YEM-1_* mutants. The different recombinant strains were grown on Terrific Broth (TB) medium supplemented with kanamycin (50 μg/mL) and chloramphenicol (30 μg/mL). The pre cultures were incubated O/N at 37°C under agitation and 40 mL were used to inoculated 1 liter of fresh TB medium supplemented as described above. IPTG (100 μM final concentration) was added when the culture reached an A^600^ of 0.7 and the cultures were incubated O/N at 18°C. Cells were harvested by centrifugation (5,000 g for 10 min at 4°C), and the pellets were resuspended in 40 mL of 50 mM MES buffer (pH 6.0; buffer A). The bacteria were disrupted at a pressure of 5,500 kPa with the help of a cell disrupter (Emulsiflex C3; Avestin GmbH, Germany). The lysates were centrifuged at 45,000 g for 30 min. The cleared supernatants were filtered in a 22 μm filter and then loaded onto a HitrapTM SP-HP 5-mL column (GE Healthcare, Belgium) equilibrated with buffer A. The enzymes were eluted with a salt gradient using buffer A and 50 mM MES, 1 M NaCl, pH 6.0 (buffer B). The fractions containing β-lactamase activity were pooled, and then dialyzed O/N in 25 mM HEPES, pH 7. The active fractions were then loaded onto a column containing pentadentate chelator (PDC - 5 mL)) equilibrated with 25 mM HEPES, pH 7.0 (buffer A). The enzymes were eluted with a salt gradient between Buffer A and 25 mM HEPES, 1M of NaCl, pH 7.0 (Buffer B). The active fractions were collected and concentrated on an YM-10 membrane (Amicon, Beverly, Mass.) to a final volume of 2 mL. Subsequently, the sample was loaded in a molecular sieve Superdex 75 GL (10/300) column (GE Healthcare, Belgium) equilibrated in 50 mM MES buffer (pH 6.0) containing 0.2 M NaCl. At the end of each purifications step, β-lactamase activity was routinely measured spectrophotometrically by following the hydrolysis of a solution of 100 μM imipenem by YEM-1.

### Kinetic characterisation

Kinetic experiments were performed by following the hydrolysis of each substrate at 30°C in 50 mM MES buffer (pH 6.0). The data were collected with a Specord 50 PLUS spectrophotometer (Analytik Jena, Germany). Each kinetic value is the mean of three different measurements; error was below 5%. Kinetic parameters were determined under initial rate and by the linearization of the Michaelis-Menten equation by the method of Hanes-Woolf. The dependence of metal ion content on activity were determined by following the hydrolysis rate of 100 μM imipenem in function of increasing ZnCl_2_, CoCl_2_ and CdCl_2_ concentration ranging from 0 to 4 mM.

### Molecular modeling

A model of YEM-1 was obtained with the Yasara software (Krieger, E., and G. Vriend. YASARA View - molecular graphics for all device - from smartphones to workstations. Bioinformatics 15:2981–2982) using the CphA structures 1X8G as templates. YEM-1 and CphA share 58% of sequence identity. The overall quality Z-score was ranked as good (−0.079), resulting from a three-dimensional packing considered good (−0.982 Z-score), and dihedrals (0.778 Z-score) and one-dimensional packing (0.679 Z-score) considered optimal. Figures were prepared using Pymol (PyMOL Molecular Graphics System, Schrödinger, LLC.)

## ACKNOWLEDGMENTS

Esposito R was supported by an ERASMUS student fellowship from the University of Federico II of Naples (Italy) and Di Costanzo S. was supported by an ERASMUS+ fellowship for traineeship from the University of L’Aquila (Italy). This work was also supported by the University of Liège and by Fund for Scientific Research (FRS-FNRS) Belgium.

We declare that there are no conflicts of interest.

---

**SUPPLEMENTAL MATERIAL** - *PS Mercuri et al*

**Table SI-.**
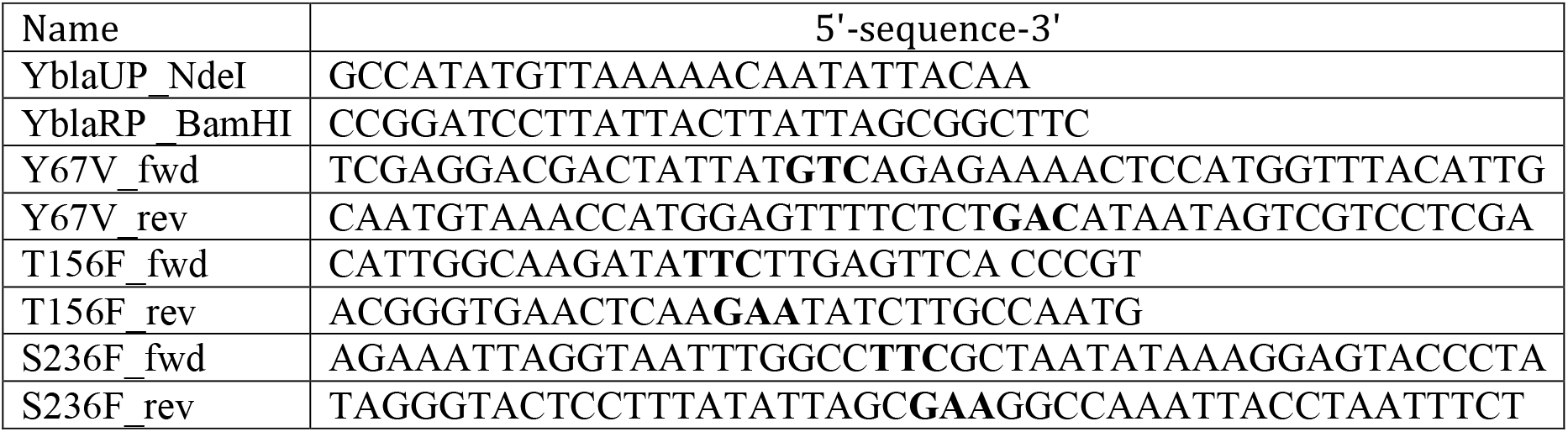
Sequence of th oligonucleotides used in this study. The mutated codons are written in bold

